# Phospholipid-independent biogenesis of a functional RP4 conjugation pilus

**DOI:** 10.1101/2025.06.27.661960

**Authors:** Naito Ishimoto, Shan He, Mikhail Bogdanov, Terry K. Smith, Gad Frankel, Konstantinos Beis

## Abstract

Conjugation, the process of DNA transfer between bacteria, is initiated universally by the formation of a mating pair formation (MPF) via a conjugative pilus. Conjugation of the IncP RP4 plasmid is mediated by short, non-retractable, rigid mating pili. Here, we report the cryo-EM structure of the RP4 pilus at 2.75 Å resolution. Uniquely, and consistently with quantitative mass spectral analysis, this revealed that the cyclic TrbC pilin subunit is not lipidated. Consistently, an *E. coli pgsA* mutant lacking phosphatidylglycerol (PG) can serve as a donor of RP4 but not of R27, encoding the H-pilus consisting of PG-associated cyclic pilin subunits (TrhA). The RP4 is the first example of a lipid-independent functional mating pilus. This discovery not only challenges the long-held assumption that an amphipathic lipid moiety is essential for the construction of conjugative pili and for MPF, but also expanding our understanding of the diverse mechanisms that bacteria employ to transfer genetic material.

## Introduction

Bacterial conjugation ^1, 2^, an efficient mechanism of horizontal gene transfer playing a central role in the dissemination of antibiotic resistance, relies on extracellular appendages known as conjugative pili (mating pili); these dynamic filaments mediate intercellular contacts between donor and recipient bacteria and facilitate DNA transfer ^3^. Studies of the archetypical F-pilus, encoded by the pOX38 and pED208 plasmids, provided detailed molecular insights into the pilus biogenesis process ^4, 5, 6^. Briefly, the TraA pilin, which is synthesized as a pre-pilin, is cleaved in the cytosol by the signal peptide peptidase LepB before it is inserted into the inner membrane (IM), where it accumulates as a reservoir. While extracted from the IM the pilin co-extracts phospholipids. The pilin-phospholipid complex then polymerizes into a helical filament on top of the type IV secretion system (T4SS), a multi-protein complex spanning the inner and outer membranes and mediating plasmid transfer ^3,4, 5, 6^. Structural studies of the pOX38/pED208/pKpQIL**-(**IncF) ^4, 7^, R388-(IncW) ^8^, and pKM101-(IncN) ^9^ encoded pili revealed that they are hollow helical tubes, approximately 8-10 nm in diameter, with a central lumen of 2.5-3 nm, where the relaxase-bound single-stranded (ss) DNA can transverse ^10^. The pilins are composed of three α-helices, α1-α3, connected by short loops; the helices display conformational variations in different pili. All known conjugative bacterial pili, the archaeal import pili ^11^ and the T-pilus encoded by the Ti-plasmid of *Agrobacterium tumefaciens* ^11, 9, 12^ have a bound phospholipid (e.g. phosphatidylglycerol – PG, phosphatidylethanolamine – PE, phospholipid GDGT-0, phosphatidylcholine – PC, and C_25_-C_25_, C_25_, C_25_ diether lipids) at a 1:1 stoichiometry with the pilin subunit. In the retractable F-pilus, the phospholipid acyl chains are parallelly arranged ^4^, while in the unretractable R388 pilus they are splayed ^8^. Cryo-electron microscopy studies have shown that each TraA pilin subunit in the pilus is associated with a single PG molecule ^4^. The PG molecule makes extensive contact with five surrounding TraA subunits, while each TraA subunit interacts with five PG molecules. Disrupting the interface between the PG and the pED208 TraA pilin by site directed mutagenesis aborted pilus biogenesis ^4^. The phospholipids act as a “molecular glue,” stabilizing the protein-protein interfaces and strengthening the entire filament. This intricate network of both protein-protein and protein-lipid interactions is crucial for assembling the helical filament. Stoichiometrically arranged PG molecules line the TraA pilin’s lumen with their solvent exposed head groups directed to the interior of the pilus and the acyl chains entirely buried between protein subunits. The negatively charged PG molecules lining the pilus lumen have been suggested to neutralize the positive charge of the pilus lumen, thus providing directionality for electronegative ssDNA transport along the pilus interior ^4^. The exception is the T pilus where the PE contributes to the positive charge within the pilus lumen ^11, 9^.

We have recently reported a deviation from the prevailing model, showing that the TrhA pilin, encoded by the prototype IncHI1 plasmid R27, undergoes posttranslational C-terminal cleavage and cyclization ^13^. Pilin cyclization is mediated via an intramolecular covalent head-to-tail peptide bond between G1 and D69, forming a macrocyclic structure. The cyclic pilin is lipidated and the overall pilus cryo-EM structure resembles the archetypical F-pilus ^4^. Like the R27 ThrA pilin, it was previously shown biochemically that the RP4 (IncP) plasmid-encoded TrbC pilin is also cyclic ^14, 15, 16^.

Several studies provided detailed molecular insights into the RP4 pilus biogenesis process ^14, 15, 16^. Following removal of the signal peptide by LepB, maturation of TrbC requires a yet unknown chromosome-encoded protease, cleaving the C-terminal 27 residues, and the plasmid-encoded TraF, which while cleaving four additional C-terminal residues cyclises TrbC by forming a peptide bond between S1 and G78 of the mature pilin ^14, 15, 16^. Here, we report the cryo-EM structure of the RP4 pilus, which unexpectedly revealed that the cyclic TrbC pilin assembles into a pilus in the absence of phospholipids. Although the cyclic structure of the pilin is thought to provide the pilus with greater stability, this discovery challenges the long-standing model of pilus formation, where an amphipathic lipid moiety was considered essential for the assembly and stability of mating pili and offers an alternative, lipid-independent pilus biogenesis pathway, that diverges from the canonical model.

## Results and discussion

### The cyclic TrbC pilin is assembled into a mating pilus in the absence of a phospholipid

We adopted the same approach to express and purify the RP4 pilus for structural studies as we reported for the H-pilus ^13^. Briefly, RP4 pili were purified from *E. coli* MG1655 Δ*fimA* (lacking expression of type I pili) using the needle-shaving and precipitation method (see Materials and Methods) (Figure 1a). Using this method, we determined the cryo-EM structure of the RP4 pilus at 2.75 Å resolution (map:map FSC, 0.143 threshold) employing helical reconstruction in cryoSPARC by imposing helical (rise 15.15 Å, twist 24.62°) and C5 symmetry (Figure 1b, c, d and Supplementary Figure 1). The outer diameter of the pilus is ~80 Å and the lumen diameter ~20 Å, comparable to other conjugation pilus structures. The Coulomb map and structure confirm that the TrbC pilin is cyclized as a result of condensation of the Ser1 amine and Gly78 carboxyl residue and agrees with the previous biochemical studies ^15^ (Figure 1c). The TrbC pilin contains three α-helices, α1-α3; the α1 and α2 helices are fused to form two α1/2 helices connected to a split α3 helix, α3_1_ and α3_2_, through a connecting loop. This arrangement resembles the conformation of the reported acyclic VirB2 pilin forming the T-pilus rather than the cyclic TrhA pilin forming the H pilus that has a split α3 helix connected via a loop to α1 and α2 helices (Figure 2a). Although the mature cyclic TrbC pilin has an additional 10 C-terminal residues compared to other pilins (Figure 2a, b), the extra residues do not impact the overall structure and can be accommodated by displacement of the α3_2_ helix by 5 degrees along Gly66 (Figure 2c). Unexpectedly, the structure revealed that the TrbC pilin is not associated with a phospholipid. Inspection of the maps did not reveal any lipid-like electron density in the proximity of the α2 or α3 helices where the bound lipids are usually located (Figure 1c, 1e, 2a). We validated that no phospholipid was present by mass spectrometry analysis on lipid extractions from the RP4 pilus (purified from cultures grown at either 25 or 37 C), and lipid extractions from the R27 pilus, which was used as a positive control. As expected R27 derived pili, showed PG species, PG 32:1 and PG 34:1, similar to those previously observed ^13^, while the lipid extract from RP4 pili showed a complete absence of lipids under two growth temperatures (Figure 3 and Supplementary Figure 2).

**Figure 1.**
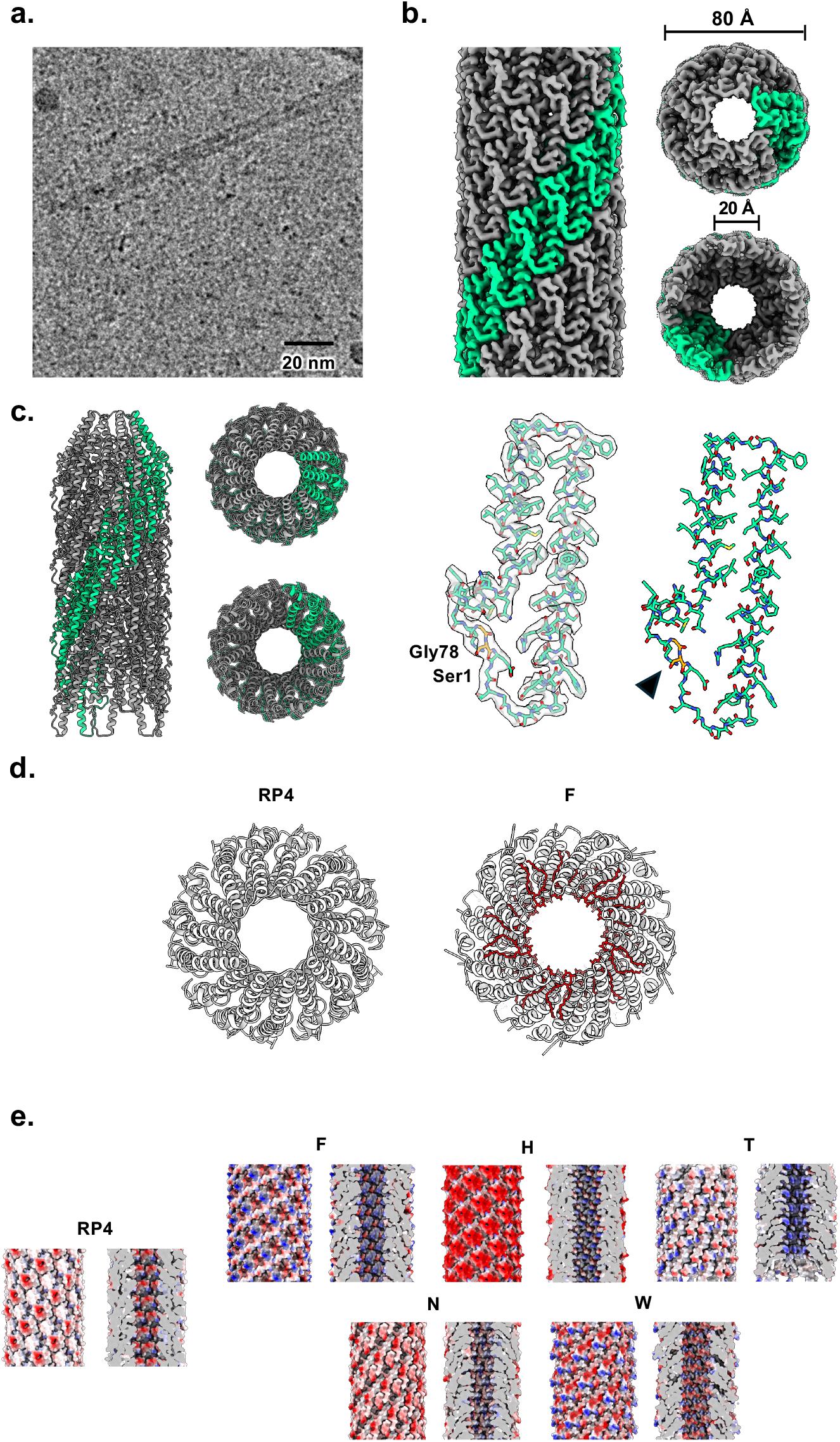
Cryo-EM structure of the RP4 pilus. (a) Micrograph of the straight and rigid RP4 pilus. (b) Cryo-EM map of RP4-pilus. One right-handed 5-start strand is shown in green and the rest of the filament in grey. The pilus has an outer diameter of ~80 Å and an inner lumen diameter of ~20 Å. (c) Atomic model of the RP4 pilus (left panel). The TrbC pilin is cyclic and lacks a bound phospholipid as revealed by the Coulomb map. The site of cyclization is indicated by an arrowhead and the residues in orange. (d) The F-pilus lumen is lined by PG (indicated by red sticks) compared to the RP4 pilus. (e) The RP4 pilus lumen displays a strong electronegative charge compared to the other bacterial pili that appear more neutral, whereas the T pilus has an alternating positive and negative charge. The exterior of the different pili displays variable charge profile.

**Figure 2.**
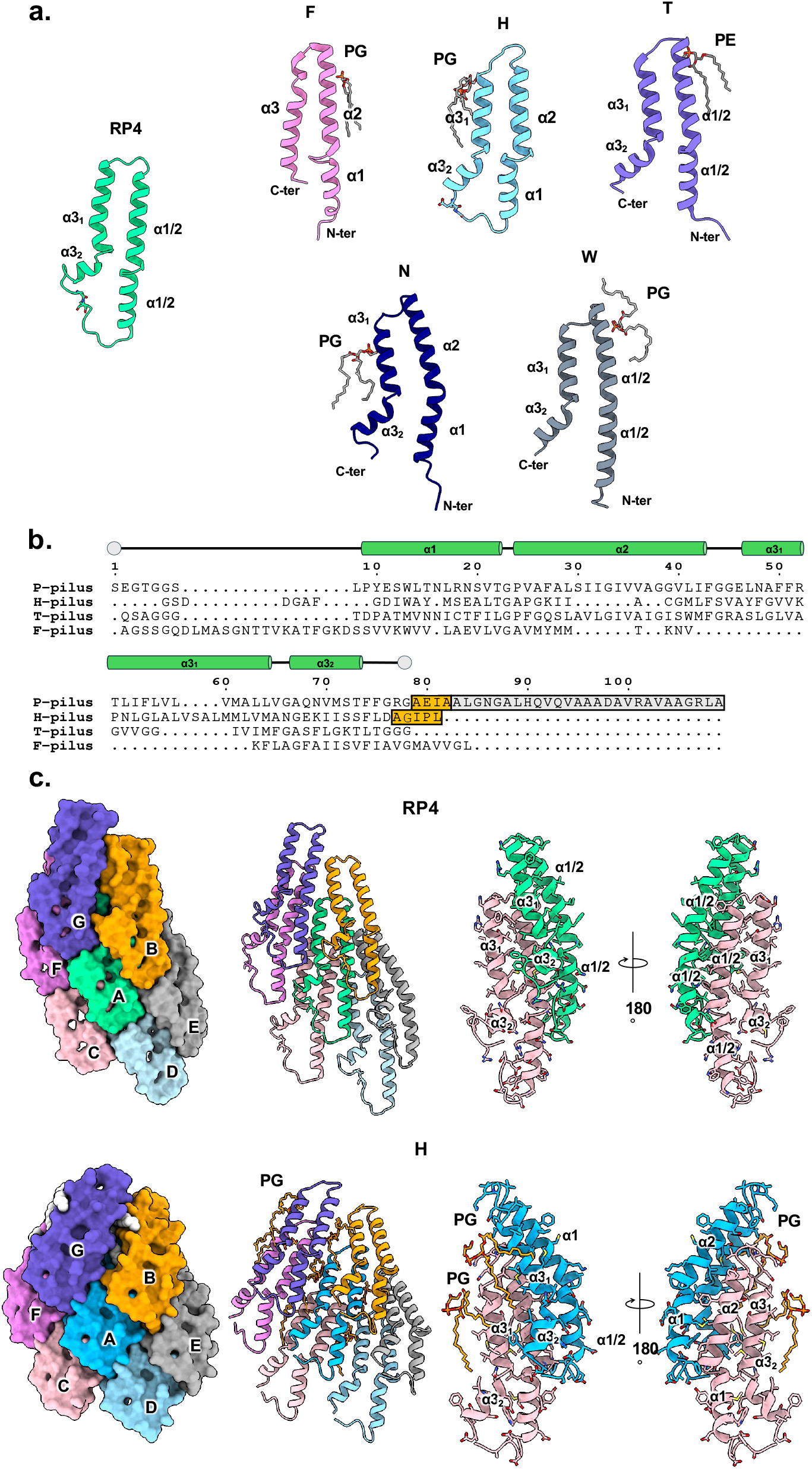
Comparison of the TrbC with other conjugative pilins. (a) The conformation of TrbC pilin resembles the VirB2. Pilins of different conjugative pili bind diverse phosholipids and in different conformation. The pilin structures and their associated lipids are shown in cartoon and sticks, respectively. (b) The C-terminal 27 residues of the pre-TrbC pilin are cleaved by an unknown chromosomal protease (highlighted in grey) and is further processed by TraF-mediated cleavage of 4 additional residues (highlighted in yellow), linked to cyclisation. The C-terminus of the F- and T-pilus that is not processed. Cyclization of the R27 ThrA pilin involves cleavage of the last 5 residues (highlighted in yellow). (c) In the absence of phospholipids, the neighbouring TrbC protomers occupy the empty lipid-space to make extensive stabilizing interactions (top row) compared to the lipidated ThrA (bottom row). Each pilin protomer is shown in either surface or cartoon representation and colored differently.

**Fig 3.**
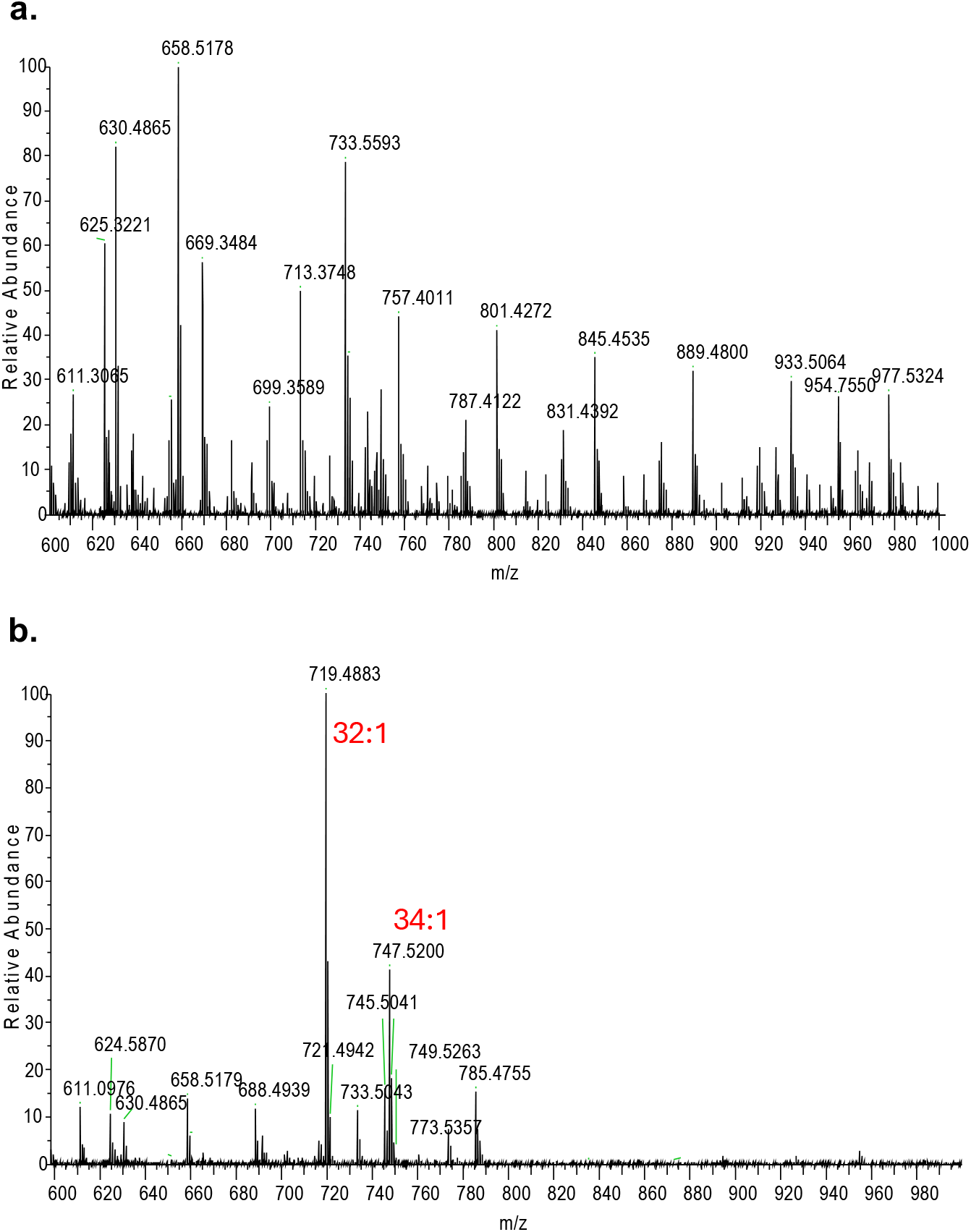
Mass spectrometric analysis of lipids associated with purified pili. Negative ion mode survey scans (600-1000 m/z) of lipids isolated from pili after treatment with phospholipase A2 (Supplementary Figure 2 shows the 100-1000 m/z). Annotated phospholipid identities were confirmed by accurate mass. (**A**) RP4 (25C) and (**B**) R27.

The lack of an associated phospholipid is unexpected as all pilus structures reported to date have been shown to be associated with inner membrane lipids needed for biogenesis. In the lipidated pilins, the phospholipid mediate contacts with neighboring pilin subunits that provide extra stability in addition to the intermolecular interactions (Figure 2c). The hydrophobic residues in TraA (pED208/F pilus) and ThrA (R27/H pilus) interact with the phospholipid acyl chain (Figure 2c). In contrast, interactions of the TrbC protomers are mediated primarily by intermolecular van der Waals forces (Figure 2c). In the absence of a bound phospholipid in the protomer interface, α3 and α2’ from the next protomer, of the TrbC protomers come closer to ‘fill-in’ the space in the absence of a phospholipid via a tight α3 and α2’ interface; this results in an offset of the two adjacent TrbC protomers by 24° rotation and 15 Å rigid body shift relative to the VirB2/ThrA protomers (Figure 2c). TrbC also makes extensive interactions with 6 surrounding subunits whereas in lipidated-pilus structures the associated phospholipids are found at the pilin interfaces (Figure 2c). The presence of different phospholipid headgroups facing the lumen of pili change their overall charge from electropositive towards to neutral/slight electronegative (Figure 1d, 1e). As the lumen of the lipid-free RP4 pilus is inherently negatively charged this could potentially negates the need for a negatively charge from the PG. The presence of a PE lipid in the T-pilus lumen results in alternating positive to negative charge; this may be because the T pilus transports diverse range of substrates beyond DNA ^11^.

### An *E. coli pgsA* mutant can serve as a donor of RP4 but not R27

To confirm the dispensable role of glycerophospholipids in RP4 pili assembly and function, we utilized a viable *pgsA* null UE54 mutant ^17, 18, 19^, completely lacking PG, as a donor. Conjugation of R27 was used as a control. The *pgsA* gene encodes phosphatidylglycerophosphate synthase, an enzyme that catalyzes the first committed step in the synthesis of PG and cardiolipin (CL). PG is a ubiquitous negatively charged phospholipid that present in almost all bacteria and performs specific head group-specific ligand functions or reactions serving as biosynthetic precursor and probably fundamental for terrestrial life ^20^. Therefore, as with the other Δ*pgsA* strains, PG-less UE54 is viable only if it carries two additional gene mutations, including an *lpp* mutation necessary to prevent accumulation of the nascent major outer membrane lipoprotein (Lpp), which requires PG for its modification. Moreover, the Rcs phosphorelay system is constitutively activated in Δ*pgsA* mutants resulting in the cell lyses at 37–42°C, which is prevented by a Δ*rcsF* mutation. The triple *pgsA lpp rcsF* mutant is viable at any temperature because it overcomes both the structural damage caused by the Lpp protein and the lethal Rcs temperature-sensitive stress response activated by the absence of PG in the *pgsA* mutant. Despite the major alterations in lipid composition UE54 cells divide and grow normally ^21^.

Although without PG and CL, the triple mutant can possesses some anionic phospholipids, such as phosphatidic acid (PA) and N-acyl-phosphatidylethanolamine (N-acyl-PE) ^21^, PA is accumulatedonly at trace amounts and N-acyl-PE is not synthesized at all in logarithmically grown cells and PE account for near 99.8% in *pgsA* null UE54 (Fig. 4A). [^32^P]PO_4_-labeling in conjunction with thin layer chromatography (TLC) demonstrates that the *ΔpgsA* mutant is producing only PE as the major zwitterionic phospholipid (Figure 4a and Supplementary Figure 3).

**Figure 4.**
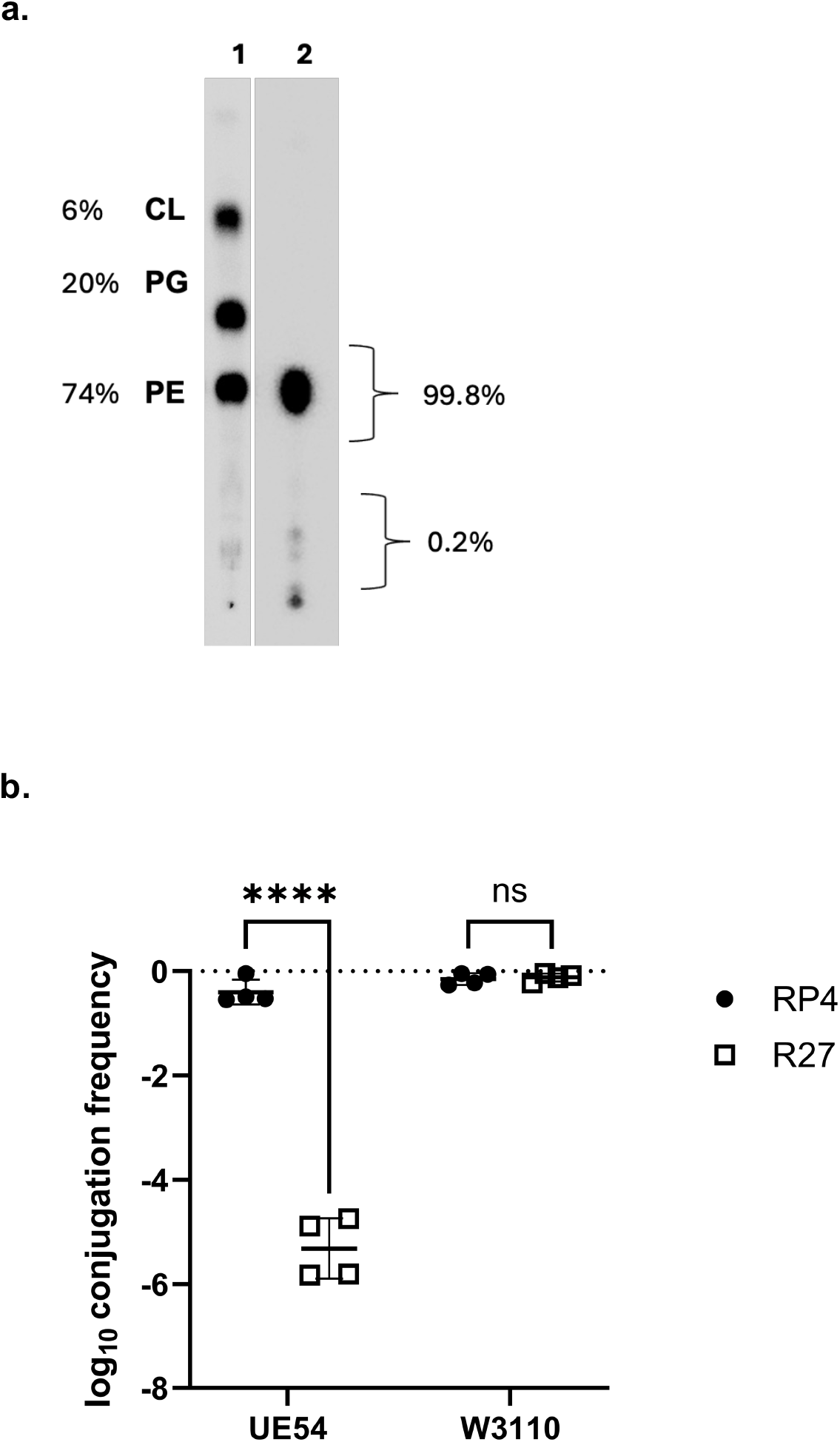
The role of PG in RP4 conjugation. A. Lipid analysis profiles of parental *E. coli* cells with normal (1) and UE54 *pgsA* null mutant (2) with altered lipid composition. The UE54 (Δ*pgsA*) contains primarily PE and no detectable PG. Uniformly ^32^P labeled phospholipids were extracted and analyzed after separation by one-dimensional TLC on boric acid-impregnated silica gel plates in solvent consisting of chloroform:methanol:acetic acid (65:25:8 v/v). B. W3110 conjugates RP4 and R27 as efficiently into an *E. coli* MG1655 recipient, while UE54 conjugates RP4 but not R27. Conjugation frequency was compared by Ordinary one-ANOVA followed by Dunnett’s multiple comparison test. ns=not significant, ***P = adjusted P < 0.0001.

Following validation, we used the *ΔpgsA* mutant, and its parental wild type strain W3110, as donors for RP4 and R27 and *E. coli* MG1655 as a recipient. While both RP4 and R27 were conjugated as efficiently from W3110, RP4 but not R27 was conjugated efficiently from the UE54 mutant (Fig. 4B). The fact that R27 plasmid cannot be conjugated efficiently when PG-less UE54 was used as a donor confirms that its pilus exhibits high selectivity for PG and presence of PG is vital for conjugation. Moreover, PG is irreplaceable/interchangeable with other negatively charged lipid (PA and N-acyl PE) present in residual amounts in the *pgsA* null strain.

The lipid-independent biogenesis of TrbC distinguishes this pilus from other conjugative pilus types, as well as from the T-pilus, which is formed from the VirB2 pilin and assembles in a fivefold helical arrangement that includes phospholipids, such as phosphatidylcholine, within its structure. The cyclic pilins in both R27 and RP4 may contribute to pilus strength under the harsh environmental conditions. However, the presence or absence of a phospholipid is not a cyclic pilin feature as the R27 pilus incorporates phospholipids into its structure at a stoichiometric ratio. The RP4 plasmid’s TrbC protein is unique among cyclic pilins because it assembles into a functional pilus without requiring a phospholipid anchor. As such, the RP4 pilus provides the first known example of a lipid-independent mating pilus, expanding our understanding of the diverse mechanisms that bacteria employ to transfer genetic material.

The distinct differences between pilus structures in bacteria (cyclic and acyclic) and archaea give fundamental insights into plasmid biology, pilus biogenesis, DNA transfer and evolution. It remains unclear why certain pili, acyclic or cyclic, such as the F- or H-pilus, adopt a 5-fold symmetry where others such as the pKpQIL pilus adopt a one-start helical symmetry ^7^; this plasticity in pilus architecture underscores the emergence of adaptable bacterial conjugation systems throughout evolution. For pili that retract, such as the F-pilus, it has been suggested that phospholipids may (i) assist in the assembly and disassembly of pilin subunits from the filament by lowering the energetic cost, (ii) provide the pilus with mechanical flexibility encountered in natural environments and (iii) repel negatively charged DNA and facilitate relaxase-ssDNA transfer. It has been suggested that although the R388 pilus is lipidated, it does not retract because of the non-canonical binding of the phosphatidylglycerol at helix α3 with its acyl chains splayed; the phospholipid would have to undergo an energetically costly rearrangement for the pilus to retract ^8^. The absence of lipid-like electron density in the TrbC maps highlights a fundamental difference in the assembly and structure of the conjugative apparatus encoded by IncP plasmids, which are found in diverse bacterial species (Supplementary Figure 4). The absence of bound phospholipids in RP4 pilus explains its rigidity and why it is not retractable ^22^. In conclusion, our data suggest that the phospholipid is not needed for TrbC extraction from the inner membrane, pilus biogenesis and relaxase-ssDNA transfer. Taken together, our finding of phospholipid-independent biogenesis redefines the role of phospholipids in conjugation pilus biogenesis and DNA transfer.

## Supporting information

Supplementary material revised

## Acknowledgements

We thank Prof Christopher Thomas, University of Birmingham, for providing us with the IncP plasmid. We thank Prof Edward Egelman, University of Virginia, for critical reading of the manuscript. We would like to acknowledge Diamond for access and support of the cryo-EM facilities at the UK national electron Bio-Imaging Centre (eBIC) under visit BI33230. This project made use the Computing Platform for Electron Microscopy at Imperial College funded by the BBSRC Mid-range equipment Initiative 22ALERT (BB/X019284/1). This work was also supported by National Institutes of General Medical Sciences Grant R01GM121493-6 to MB. NI is funded by a Fellowship from The Naito Foundation. GF is supported by a Wellcome Trust Investigator Award grants 224282/Z/21/Z.

## Author contributions

KB and GF conceived and supervised the study. NI purified the RP4 pilus, determined the cryo-EM structure and performed model building and refinement. SH conducted conjugation assays. MK generated and analyzed the Δ*pgsA* mutant. TKS performed the lipidomics assays NI, GF and KB analyzed the structures. NI, MK, GF and KB wrote the manuscript.

## Materials and Methods

### Expression and purification of RP4 pilus

The RP4 pilus was purified following the protocol used to prepare the H pilus ^13^. Briefy, bacterial cells from an overnight culture were shaved through 25 G needles 25~30 times and were centrifuged at 50,000 xg for 1 h. The pilus filaments were precipitated using PEG 6,000 at a final concentration of 5% with stirring for 1 h, which was followed by centrifugation at 50,000 xg for 30 min. The supernatant was removed, and the pellet was resuspended in 200 μL imaging buffer (50 mM Tris-HCL pH 8, 200 mM NaCl). The solution containing the filaments was dialyzed overnight in imaging buffer. Sample purity was judged by SDS-PAGE.

### Grid preparation and cryo-EM data collection

For grid preparation, 3 μL of the purified sample at ~4 mg/mL was applied to a glow-discharged carbon grid (Quantifoil R1.2/1.3, Cu, 300 mesh). The grid was blotted with a blotting force of 0 for 3 sec at 4°C and 100% humidity, and flash-frozen in liquid ethane using a Vitrobot Mark IV (Thermo Fisher Scientific). Cryo-EM data were collected on 300 kV Titan Krios G2 (Thermo Fisher Scientific) with a Falcon 4i detector and Selectris X energy filter at the Electron Bioimaging Center (eBIC), UK. A total of 3,405 movies were collected at a magnification of ×130,000 (pixel size of 0.921 Å/pixel) with a total electron dose of 50 e/Å2. The defocus range was set between −0.8 and −1.8 μm. All data were automatically acquired using EPU software. The data collection parameters are summarized in Supplementary Table 1.

### Cryo-EM data processing and model building

The collected data were processed using cryoSPARC (v.4.6.1) ^23^. Data were collected from a Titan Glacios and Krios, analyzed separately, and 2D images and initial volume from Glacios dataset were used for initial reconstruction. Both datasets were processed following similar steps. Motion correction was performed using Patch Motion Correction, and CTF estimation for micrographs was done by Patch CTF estimation ^23^. Particles from the Glacios dataset were manually picked from all micrographs and extracted in a binned state at 2.76 Å/pix. The extracted particles were classified in two rounds in 2D classification, and selected high-quality particles were used as references for the Filament tracer. Three rounds of 2D classification from the whole dataset was used to select particles for helical reconstruction. Helical symmetry search job suggested an axial rise of ~15.3 Å and a twist of ~24.6°, along with 5-fold point group symmetry. These symmetry parameters were used as starting parameters for helical reconstruction using Helix refine. 4.0 Å P-pilus map was reconstructed from Glacios data set. Then, 2D references and 3D model were used for reference to Krios dataset. In Krios dataset, 3,405 movies were collected. After Motion Correction and CTF refinement, template pick was performed with Filament tracer using a 2D reference from the Glacios dataset. 4,302,764 particles were extracted in the binning state. Three rounds of 2D classification were performed, and 10,792 particles were selected. Using 3D volume and helical parameters from the Glacios dataset, Helical refinement, local CTF refinement and Reference-based motion correction were performed several times. The final RP4 pilus helical reconstruction showed 2.75 Å resolution using 10,729 particles, with a rise of 15.15 Å and a twist of 24.62°.

*Ab initio* model building and real space refinement were performed in PHENIX ^24, 25^. The model was subjected to iterative cycles of refinement and manual rebuilding in COOT ^26^. The structural model was validated using MolProbity ^27^. The processing parameters and refinement statistics are summarized in Supplementary Table 1.

### Determination of steady-state phospholipid composition by radiolabeling

To determine the steady state phospholipid composition by radiolabeling, cells were uniformly labeled with 5 μCi/ml of [^32^P]PO4 after dilution to OD_600_ of 0.05 to initiate logarithmic growth after dilution of the overnight culture. Cells were harvested by centrifugation, and phospholipids were extracted by acidic Bligh Dyer procedure. After labeling, cells were harvested by centrifugation and resuspended in 0.2 mL of acidic solution (0.5 M NaCl in 0.1 N HCl). Lipids were extracted by adding 0.6 mL of solution 2 (chloroform:methanol, 1:2, v/v), followed by vortexing for 30 minutes. Subsequently, 0.2 mL of resuspension buffer was added, and the sample was vortexed for an additional 10 minutes. Phase separation was achieved by centrifugation at 13,000 × *g* for 5 minutes. The lower chloroform phase, containing the extracted phospholipids, was collected. An aliquot corresponding to approximately 1,000– 2,000 counts per minute (cpm) of radiolabeled phospholipid was used for thin-layer chromatography (TLC) analysis. One-dimensional TLC was performed to resolve individual phospholipid species. Boric acid-impregnated Merck Kieselgel 60 TLC plates (0.25 mm thickness) (EMD, Gibbstown, NJ) were activated and developed with a solvent consisting of chloroform:methanol:acetic acid (65:25:8 v/v). The spots were assigned to different *E. coli* phospholipids based on TLC mobility (Rf values) of in-house standards and utilization of different lipid mutants including viable *pssA* null and therefore not synthesizing major phospholipid PE. Radiolabeled lipids were detected using a Typhoon FLA 9500 PhosphoImager (GE) and quantified with ImageQuant™ software. Stored images were processed and quantified using ImageQuantTM software. Phospholipid content is expressed as mol% of total phospholipid based on the intensity of the captured signal on the Phosphor screen generated by the radiolabeled spots on the TLC plate. The results presented are representative of three or more determinations. All TLC analyses were performed in at least two independent experiments to ensure reproducibility.

### Lipid analysis of Pili by mass spectrometry

Purified pili were subjected to lipid extractions as previously described ^4^. Briefly purified pili (~0.05 mg) were treated with phospholipase A2 (0.1 units) in PBS for 16 h at 37°C, followed by heat inactivation. The remaining Bligh and Dyer extracted lipids were analyzed with a Thermo Exactive Orbitrap mass spectrometer both positive and negative ion modes.

### Bacterial strains, growth conditions and Conjugation

UE54 (*lpp*-2 Δ*ara714 rcsF*::mini-*Tn10 cam pgsA*::FRT-*kan*-FRT) is a null for the *pgsA* gene and is therefore unable to synthesize phosphatidylglycerol (PG). W3110 or UE53 (MG1655 *lpp*-2 Δ*ara714 rcsF*::mini-*Tn10* cam) were utilized as parental strains with wild-type phospholipid composition. Cells were grown in LB medium containing 10 g/L Bacto-tryptone, 5 g/L Bacto-yeast extract, and 10 g/L NaCl.

*E. coli* W3110 or UE54 containing the derepressed R27 or RP4 were used as donors (both plasmids encode chloramphenicol resistance); Streptomycin (Strp) resistant *E. coli* MG1655 was used as a recipient. The donor and recipients were grown in LB at 37 °C overnight, mixed in a ratio of 10:1 (R27) or 1:1 (RP4), diluted 1 in 25 in PBS and 40 μL spotted onto LB agar plates (no selection). Following incubation at 37 °C for 3 h, R27 conjugation was allowed to proceed overnight at 25 °C. For RP4 conjugation the mixture was incubated overnight at 30 °C. The conjugation mixtures were collected and resuspended in 1 mL of PBS following serial dilution from 10^−1^ to 10^−8^. For selection of transconjugants, the serial dilutions were plated on LB containing 30 μg/mL chloramphenicol and 50 μg/mLStrp and incubating at 37 °C overnight. Data are presented as proportions of transconjugants/recipients.

## Data availability

The cryo-EM maps have been deposited in the Electron Microscopy Data Bank (EMDB) under accession code EMD-65024. The structural coordinates have been deposited to the RCSB Protein Data Bank (PDB) under the accession code 9VF4. Raw images were submitted to the Electron Microscopy Public Image Archive (EMPIAR) with ID EMPIAR-XXXX.

