## Supplementary material revised for "Phospholipid-independent biogenesis of a functional RP4 conjugation pilus"

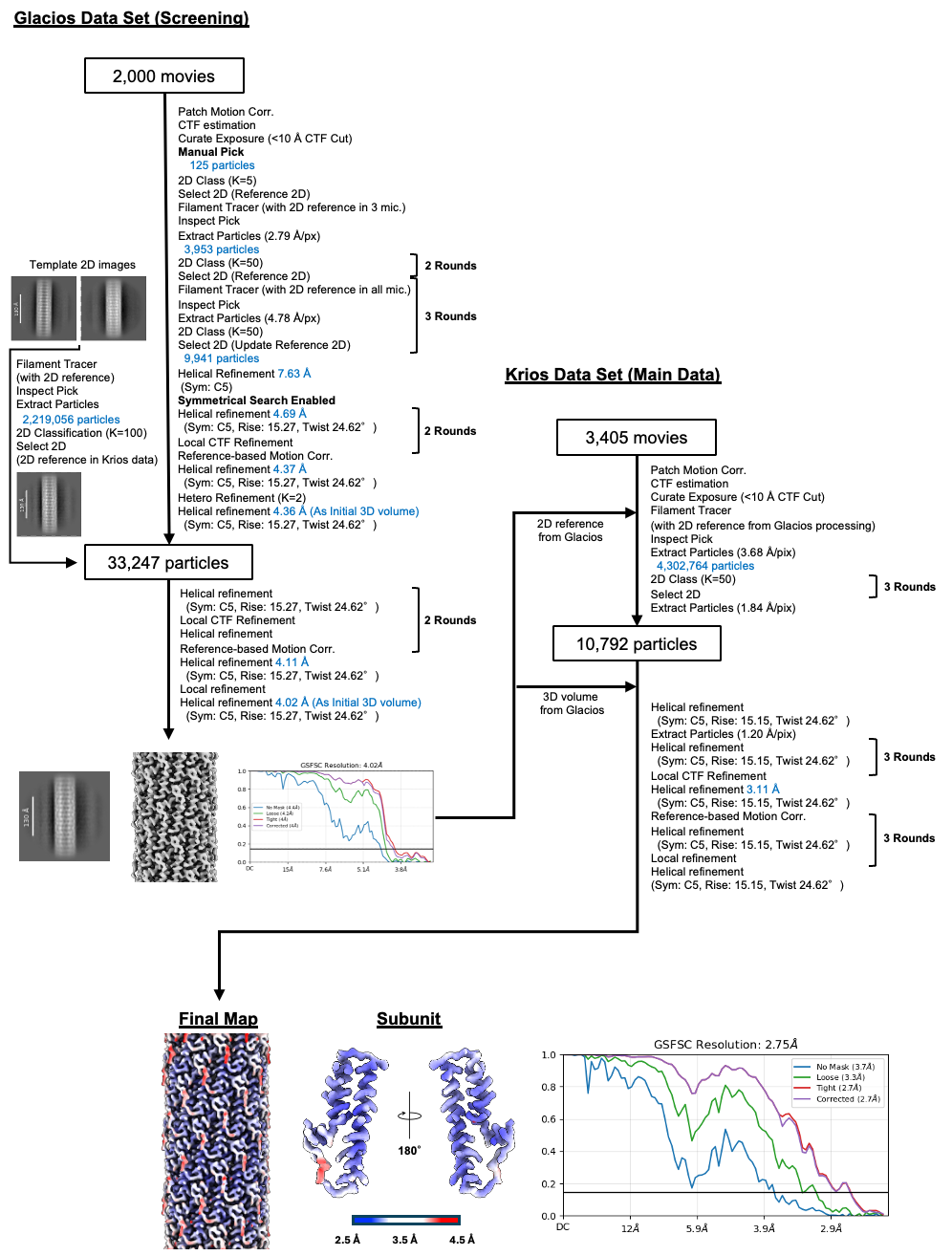

**Supplementary Figure 1.** Cryo-EM data processing. All the processing was performed in cryoSPARC (v.4.6.1). The final map was colored according to the local resolution. Gold-standard FSC curve of the final map was shown. The resolution was cut-off at FSC=0.143.

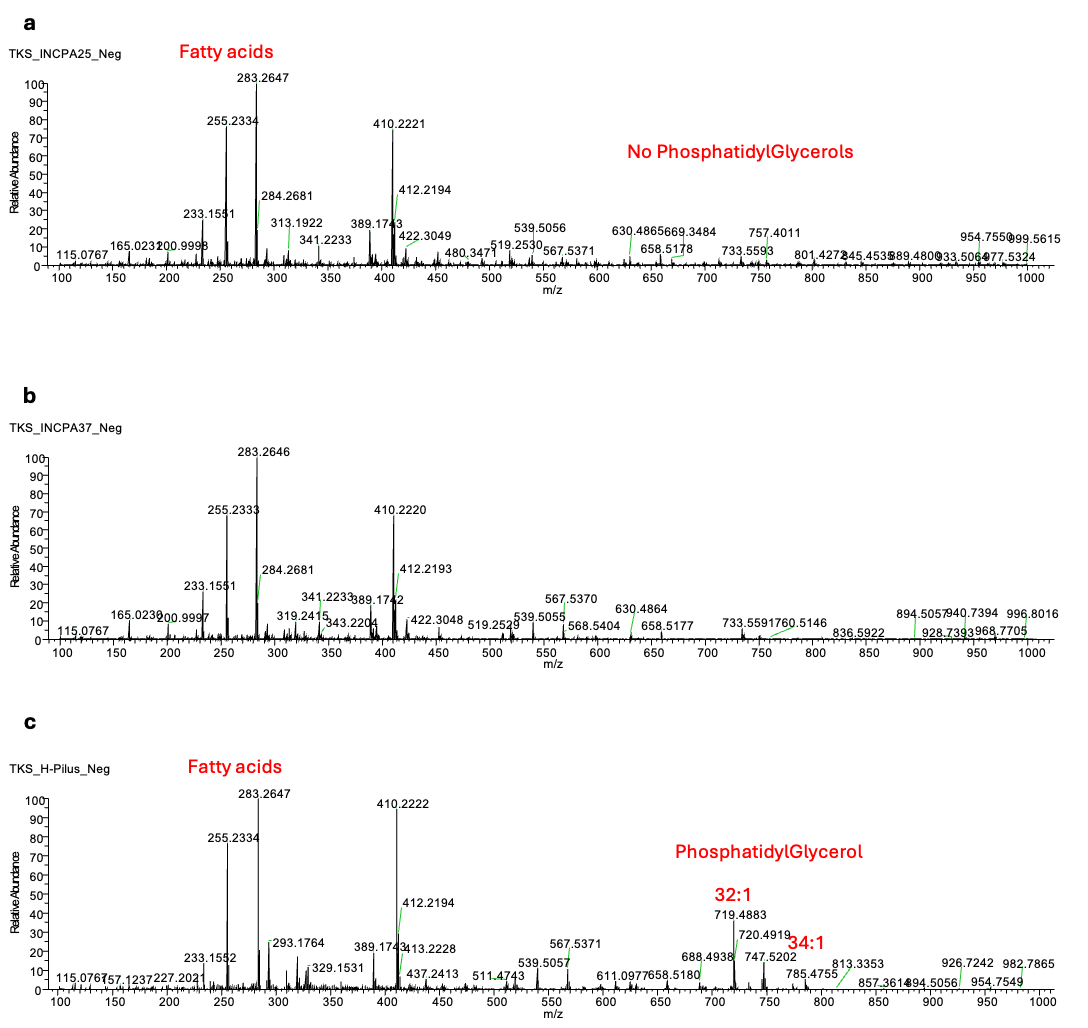

**Supplementary Figure 2.** Mass spectrometric analysis of lipids associated with purified pili. Negative ion mode survey scans (100-1000 m/z) of lipids isolated from pili after treatment with phospholipase A2. Annotated phospholipid identities were confirmed by accurate mass. (**A**) RP4 (25C); (**B**) RP4 (37C) and (**C**) R27

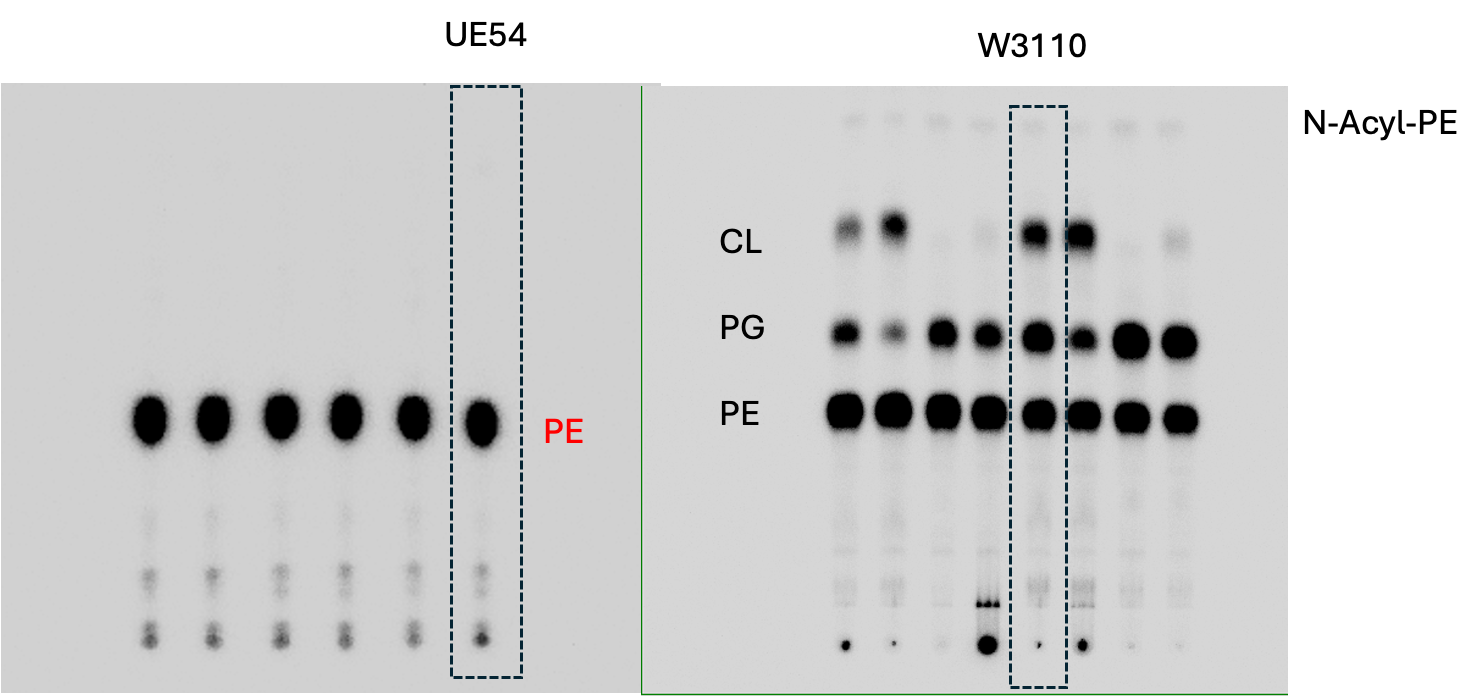

**Supplementary Figure 3.** Uncropped gels from Figure 4. Dashed lines indicate the lanes used in Figure 4.

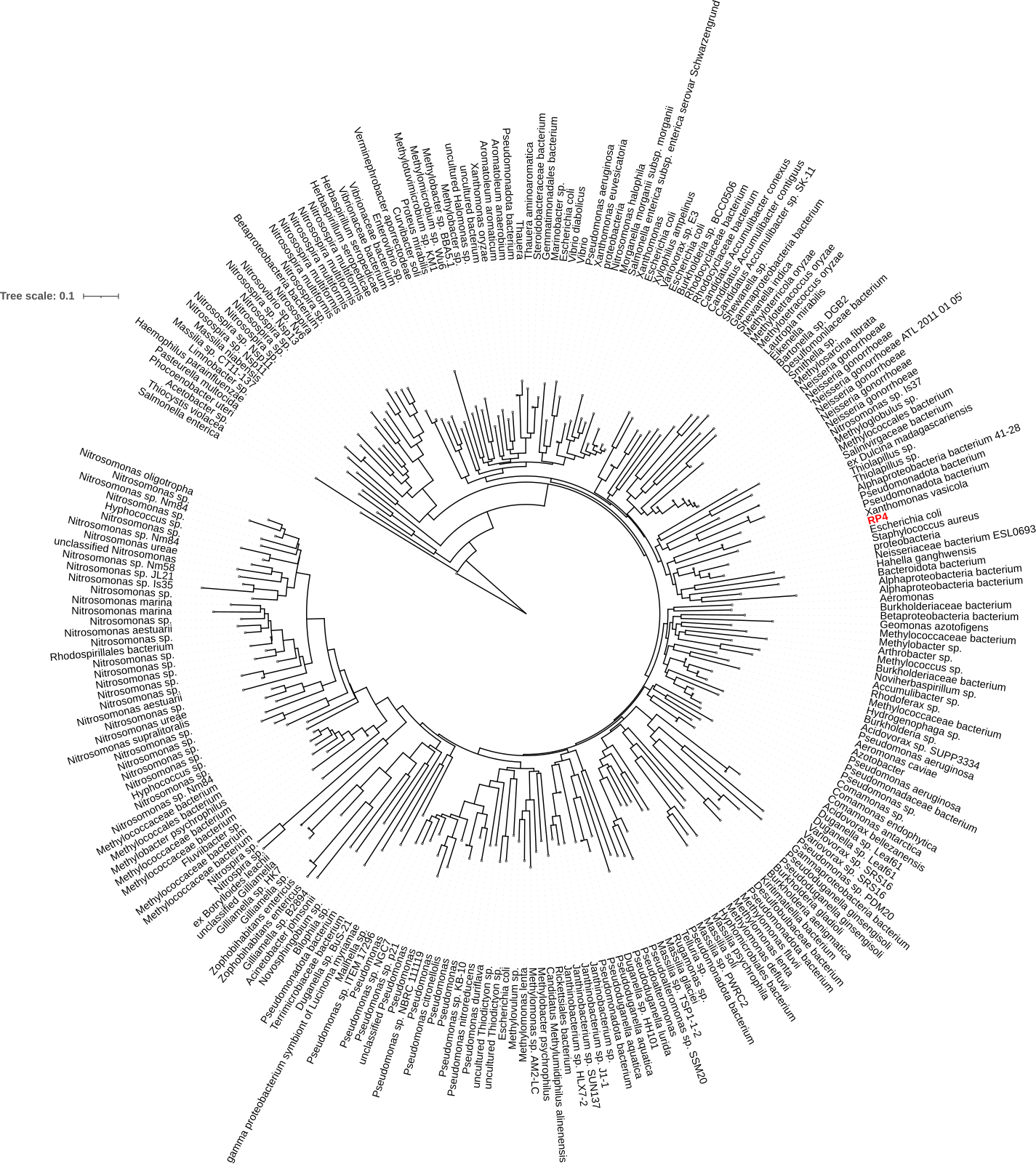

**Supplementary Figure 4.** Phylogenetic analysis of TrbC pilin. Circular phylogenetic tree at level of bacterial species. The tree was generated with NCBI BLASTP and viewed in ITOL.

**Supplementary Table 1.** Cryo-EM data collection and refinement statistics

|  | P-pilus |
| --- | --- |
| **PDB entry** | 9VF4 |
| **EMDB entry** | EMD-65024 |
| **Data collection and processing** | |
| Magnification | 130,000 |
| Microscope | Titan Krios |
| Voltage (kV) | 300 |
| Detector | Falcon 4i |
| Electric exposure (e^-^/Å) | 50 |
| Deforcus range (µm) | -0.8 to -1.8 |
| Pixel size (Å) | 0.921 |
| Data Processing Program | cryoSPARC (v.4.6.1) |
| Movies | 3,405 |
| Initial / Final particle images (no.) | 492,091 / 123,942 |
| Symmetry imposed | C5 |
| Helical rise (Å) | 15.15 |
| Helical twist (°) | 24.62 |
| Map resolution (Å) | 2.74 |
| FSC threshold | 0.143 |
| **Refinement** |  |
| Refinement Program | PHENIX (v.1.20.1) |
| Model resolution (Å) |  |
| FSC threshold = 0.143 | 2.75 |
| Model composition |  |
| Non-hydrogen atoms | 574 |
| Protein residues | 78 |
| R.m.s. deviations |  |
| Bond length (Å) | 0.006 |
| Bond angles (°) | 0.503 |
| Validation |  |
| MolProbity score | 1.38 |
| Clashscore | 4.29 |
| Ramachandran plot |  |
| Favored / Allowed (%) | 98.68 / 1.32 |
| Disallowed (%) | 0.00 |
| Mask CC | 0.86 |
